# Test-retest reliability of sensorimotor activity measured with spinal cord fMRI

**DOI:** 10.1101/2025.09.07.674708

**Authors:** Olivia S. Kowalczyk, Sonia Medina, Alessandra Venezia, Dimitra Tsivaka, Aminul I. Ahmed, Steven C. R. Williams, Jonathan C. W. Brooks, David J. Lythgoe, Matthew A. Howard

## Abstract

Establishing the reliability of spinal cord functional magnetic resonance imaging (fMRI) is critical before employing it to assess experimental or clinical interventions. Previous studies have mapped human motor activity primarily to the ipsilateral ventral horn, aligning with myotomal and dermatomal projections. Despite these insights, the test-retest reliability of spinal fMRI remains under-investigated. Here we assessed spinal cord activation during a sensorimotor paradigm involving right-hand grasping and grip force estimation in 30 healthy volunteers. Participants completed two identical scanning visits, each time performing the same task twice, enabling the investigation of test-retest reliability both within a single experimental visit and between visits performed on different days. Aggregating all task runs, motor-evoked activation was observed in ipsilateral ventro-dorsal regions of spinal segmental levels C5-T1, as well as in medial regions of levels C2-C3. Despite highly reliable task performance (grip force) and fMRI signal quality (temporal signal-to-noise ratio), the reliability of motor activation was predominantly *poor*-to-*fair* both within and between visits, with notable variability in spatial distribution observed across task runs. Increasing the number of task runs per individual improved the robustness of group-level activation, as indexed by higher activated voxel count, larger cluster spatial extent, and attenuated t-statistic distribution. Although we demonstrated that motor-evoked activation corresponds to the known neuroanatomical organisation of motor circuits, its low test-retest reliability presents a challenge for wider applications of spinal fMRI. Understanding the drivers of low reliability in functional imaging is warranted, but we suggest that looking beyond measurement error is required, including careful consideration of inherent within-individual variability underpinned by neurophysiological and psychological factors.

## 1 Introduction

The spinal cord is an integral part of the motor system, playing a key role in movement execution and control. While the function of supraspinal elements of the motor system has been extensively characterised using imaging, less focus has been devoted to the study of the spinal cord contributions to human motor function (Lemon, 2008; Nielsen, 2016; Rowe & Siebner, 2012). Recent technological and analytical developments have optimised image acquisition (Barry et al., 2021; Chu et al., 2023; Islam et al., 2019; Tsivaka et al., 2023) and processing (Banerjee et al., 2025; Brooks et al., 2008; De Leener et al., 2017), allowing non-invasive study of the spinal cord using functional magnetic resonance imaging (fMRI). Since the early applications of spinal fMRI (Yoshizawa et al., 1996), a growing literature has assessed spinal cord motor function and somatosensation, as well as the presence of spatiotemporally organised intrinsic activity of the cord, i.e. resting states (for reviews see Kaptan et al., 2024; Kinany et al., 2022; Landelle et al., 2021).

In the study of sensorimotor function, the field has employed several paradigms, primarily focussed on the upper limbs, including hand grasping, finger tapping, and extension-adduction-abduction movements of isolated parts of the limb (Landelle et al., 2021). Across paradigms, unilateral motor activity in healthy adults has been shown to elicit activation predominantly in ipsilateral ventral (motor neuron-rich) regions of spinal segmental levels corresponding to the myotomal projections of the moving body part (Barry et al., 2021; Braaß et al., 2023; Hemmerling et al., 2023; Landelle et al., 2021), aligning with the known functional organisation of the cord (Lemon, 2008; Nielsen, 2016). However, motor-related activity has also been observed in contralateral and dorsal (somatosensory neuron-rich) regions of the same spinal segmental level, sometimes with rostrocaudal distribution spanning neighbouring or more distal segmental levels (Hemmerling et al., 2023; Landelle et al., 2021). Additionally, reports indicate that the magnitude of spinal cord activation is modulated by the force generated (Braaß et al., 2023; Madi et al., 2001; Oliva et al., 2025), movement complexity (Maieron et al., 2007; Oliva et al., 2025, 2025; Vahdat et al., 2015), and motor learning (Khatibi et al., 2022; Kinany et al., 2023; Vahdat et al., 2015). Together, this literature demonstrates that spinal fMRI is sensitive to detecting not only gross motor-related activity but also more intricate processes facilitating sensorimotor function, which may relate to early processes of integration, proprioception, and motoneuron facilitation-inhibition (Landelle et al., 2021).

The findings from studies in healthy individuals suggest that spinal fMRI might be a promising clinical tool, potentially facilitating identification of disease biomarkers or predictors of treatment progression in conditions affecting sensorimotor pathways, including multiple sclerosis, spinal cord injury, and myelopathy, among others (Kinany et al., 2022; Landelle et al., 2021; Powers et al., 2018). To facilitate this clinical translation, spinal fMRI still needs to demonstrate reliable and reproducible outcomes. This is particularly important given that acquiring fMRI recordings from the spinal cord is uniquely complicated by the cord’s anatomy and location in the body. The small diameter and large rostrocaudal extent of the cord, its proximity to tissues of differing magnetic susceptibility and sources of physiological noise limit the quality of spinal fMRI data and can directly impact the reliability of techniques used to assess function (Kinany et al., 2022). Despite these important considerations, the test-retest reliability of the spinal fMRI remains under-investigated.

To date, test-retest reliability of spinal fMRI has been primarily assessed in resting-state paradigms (Barry et al., 2016; Hu et al., 2018; Kaptan et al., 2022; Kowalczyk et al., 2024; Liu et al., 2016), where reliability estimates were shown to be mixed but comparable to those of brain fMRI (Noble et al., 2019). Only two studies investigated test-retest reliability of motor-related spinal cord activation (Bouwman et al., 2008; Weber et al., 2016b), showing *good* reliability (intraclass correlation coefficient, ICC = 0.8) of average percentage signal changes in neuroanatomically relevant sections of the cord (Bouwman et al., 2008) but *poor* reliability (ICC range = 0.1-0.3) when assessed voxel-by-voxel (Weber et al., 2016b). Although these two studies offer early evidence of test-retest reliability of motor signalling recorded by spinal fMRI, reliability was only assessed within the same experimental session and in small samples (N = 3 and N = 11). Establishing the reliability of spinal fMRI measurements across distinct sessions and in well-powered samples is critical before employing it to assess experimental or clinical interventions.

The aims of this study were two-fold: (1) we investigated cervical spinal activation during a unilateral sensorimotor paradigm, involving right hand grasping (squeezing) with grip force measurement in a well-powered sample of healthy adults (N = 30, tested four times); (2) we characterised test-retest reliability of the acquired signal both within the same testing visit and across visits conducted on separate days. To do so, we used our locally-developed blood oxygen level dependent (BOLD) sensitive echo planar imaging (EPI) sequence, optimised for cervical spinal imaging and previously shown to provide increases in signal-to-noise ratio (tSNR) and a marked reduction in signal dropout (Kowalczyk et al., 2024; Tsivaka et al., 2023).

Our preregistered hypotheses were (Kowalczyk et al., 2021):

1. Task-related BOLD responses will be observed in the ipsilateral dorsal and ventral spinal cord in the C6-C8 spinal segmental levels.
2. Spinal cord responses will be reliable, with ICC test-retest reliability statistics greater than 0.4.

## 2 Material & methods

### 2.1 Participants

Data from 30 healthy right-handed (assessed using the Edinburgh Handedness Inventory; Oldfield, 1971) adult volunteers (17 females, 13 males; 19-38 years old, mean ± SD = 24 ± 5 years) survived data quality assurance and were retained for analysis. Full inclusion and exclusion criteria for this study are outlined in the study preregistration (Kowalczyk et al., 2021). Briefly, participants were excluded due to: (1) history of psychiatric, medical, or psychological conditions, (2) history of substance or alcohol abuse, (3) regular use of medications affecting the central nervous system (CNS), (4) irregular menstrual cycle for menstruating participants, (5) MRI-related contraindications. Additionally, participants were excluded if they were unwilling to adhere to the following lifestyle guidelines before each visit: (1) abstain from alcohol for 24 hours, (2) limit consumption of caffeinated drinks to one on each study day, (3) abstain from using non-steroidal anti-inflammatory drugs (NSAIDs) or paracetamol for 12 hours, (4) abstain from nicotine-containing products for 4 hours. Additional exclusion criteria based on data quality were used. Subjects with incomplete functional acquisition, poor functional image contrast, and large geometric distortions were excluded. Full details of participant/data exclusions at each stage of the study are shown in Figure 1.

**Figure 1.**
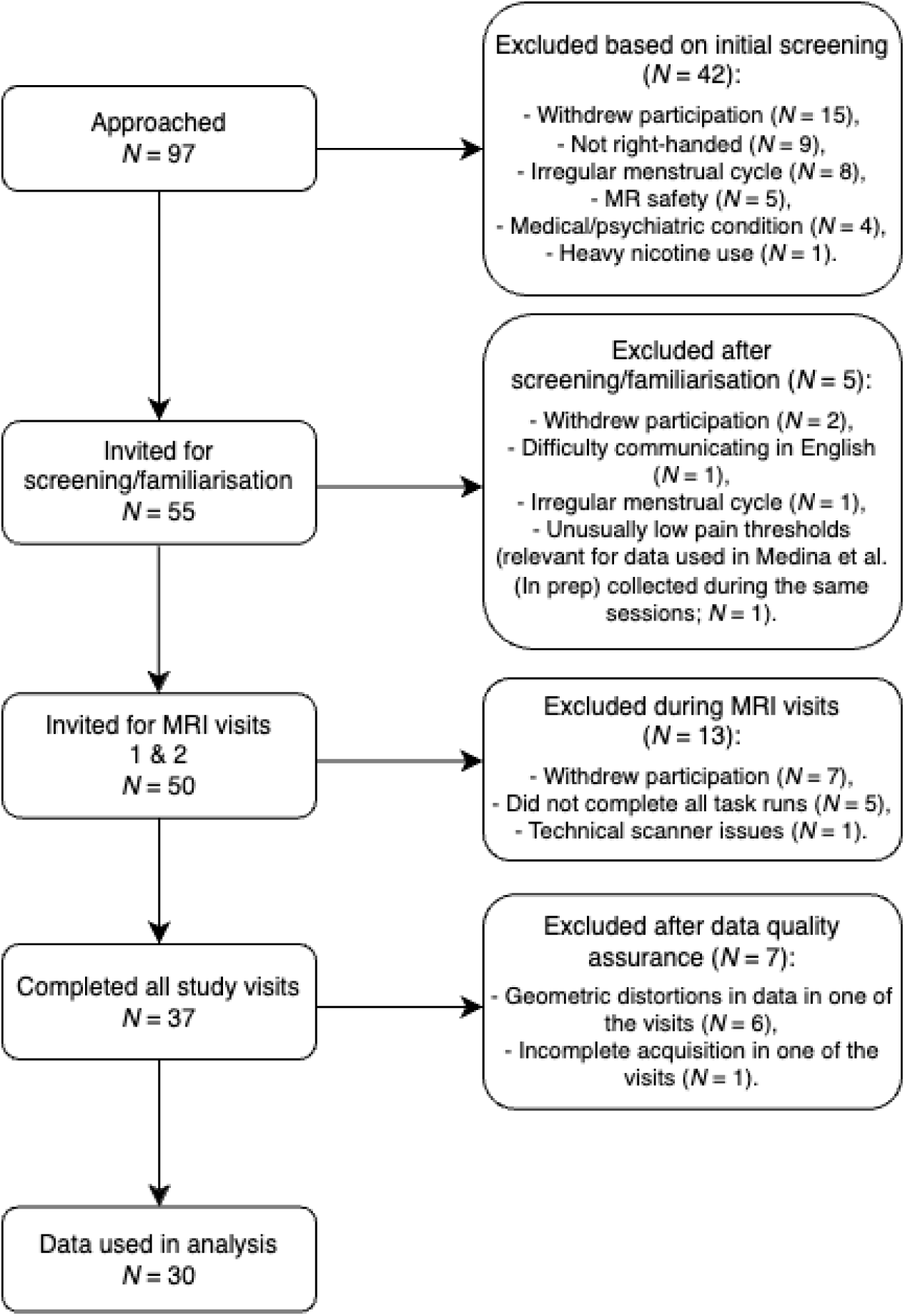
Selection of participants fulfilling the study eligibility criteria and data quality assurance. MR, magnetic resonance; MRI, magnetic resonance imaging.

Written informed consent was obtained from all participants. This study was approved by the Psychiatry, Nursing, and Midwifery Research Ethics subcommittee at King’s College London, UK (HR-16/17-4769) and conducted in accordance with the Declaration of Helsinki.

### 2.2 Procedure

The study comprised three visits – a screening/familiarisation visit and two identical MRI visits (for test-retest assessment). Across all three study visits, the mean interval (± SD, range) was 21 (± 20, 1-84) days. The mean (± SD, range) interval between the two scanning visits was 27 (± 24, 1-84) days. Variability in between-scan intervals was driven by considerations regarding diurnal (Webb et al., 2024) and menstrual cycle (Schuster & Jansen, 2022) influences on test-retest estimates, as well as practical constraints relating to scanner/participant availability and COVID-19 isolation rules in place at the time of data collection (October 2021 – March 2023). The wider study involved additional fMRI assessments of resting-state (Kowalczyk et al., 2024) and evoked pressure pain, which preceded the sensorimotor task described here. Details of those can be found in the study preregistration (Kowalczyk et al., 2021).

#### 2.2.1 Screening and familiarisation visit

Compliance with study lifestyle guidelines (see Section 2.1) was assessed at the beginning of the visit. Participants underwent breath alcohol and urine drugs of abuse tests to check alcohol/substance use. Caffeine, nicotine, and NSAIDs/paracetamol intake were assessed by self-report. Participants were familiarised with the scanner environment by visiting a mock scanner.

#### 2.2.2 MRI visits

Both MRI visits were performed at the same time of day and the same phase of the menstrual cycle for menstruating participants. The visits began with an assessment of compliance with the study lifestyle guidelines. Additionally, participants completed the state version of the State Trait Anxiety Inventory (STAI; Spielberger et al., 1971) to assess differences in anxiety levels between visits. No differences were observed (t(29) = 1.80, *p* = 0.082, Cohen’s d = 0.33, 95% CI [-0.69; 0.04]; visit 1 mean ± SD = 28.43 ± 6.35; visit 2 mean ± SD = 30.30 ± 7.92).

Participants were positioned in the scanner supine, with foam padding around the head and neck to minimise movement and neck curvature. MRI data were collected in the following order: (1) optimisation of static 0^th^, 1^st^, and 2^nd^ order shims and linear slice-specific shims, (2) structural data acquisition, (3) 10 min 50 s resting-state scan (reported in Kowalczyk et al., 2024), (4) evoked pressure pain task (Kowalczyk et al., 2021), and (5) two identical runs of a hand grasping sensorimotor task. Respiratory and cardiac traces were recorded with respiratory bellows and a pulse oximeter respectively, along with scanner triggers (at the start of each repetition time; TR), throughout the scan.

The hand grasping task required participants to squeeze a rubberised air-filled ball with their right (dominant) hand in response to verbal cues displayed on a projector screen. The screen was positioned at the rear of the scanner bore and visible through a mirror mounted on the head coil. The task was an A/B (squeeze/rest) design comprising eight squeeze/rest block pairs. Each block contained either eight ‘squeeze and release’ or eight ‘rest’ trials (trial duration = 2.5 s; block duration = 20 s). Participants completed two consecutive 5 min 20 s runs of the task per visit, resulting in a total of four task runs over the course of the study. Each run was preceded by individual hand-grip force thresholding, asking the participant to squeeze the ball as hard as they could three times. Grip force exerted was recorded throughout the task.

### 2.3 MRI acquisition

Data were acquired at the NIHR Wellcome King’s Clinical Research Facility, King’s College London using a 3 T GE MR750 System (General Electric, Chicago, Illinois) equipped with a 12-channel head-neck-spine coil and a 4-channel neurovascular array. A sagittal 3D CUBE T2-weighted structural image was acquired at the beginning of the scanning session over 64 slices with a coverage of the whole brain and cervical spinal cord to vertebral level T1 (TR = 2.5 s, echo time (TE) = 120 ms, echo train length = 78, flip angle = 90°, field of view (FOV) = 300 mm, acquisition matrix = 320 × 320, slice thickness = 0.8 mm). This acquisition was based on Cohen-Adad et al. (2021) with the FOV increased to 300 mm.

Static 0^th^, 1^st^, and 2^nd^ order shims were optimised prior to functional data acquisition. A spectral-spatial excitation pulse was used to excite only tissue water and slice-specific linear shims were implemented by adding 0.6 ms duration x, y, and z-gradient lobes after the excitation pulse. Second-order shimming and x, y, and z-shimming were optimised over elliptical regions of interest (ROIs) covering the cord drawn manually by the researcher present during scanning (OSK or SM). To maintain consistency and avoid potential systematic differences in ROI drawing affecting test-retest estimates, the same researcher drew ROIs for both MRI visits within participants. Full details of shimming optimisation for this acquisition sequence can be found in Tsivaka et al. (2023).

Functional EPI data were acquired over 38 sequential slices in descending order (slice thickness = 4 mm, slice gap = 1 mm), with the inferior-most slices anchored at vertebral level T1 and covering the whole cervical spinal cord and the brainstem (TR = 2.5 s, TE = 30 ms, flip angle = 90°, ASSET factor = 2, FOV = 180 mm, acquisition matrix = 96 × 96, reconstruction matrix = 128 × 128, in-plane voxel size = 1.41 × 1.41 mm). Four dummy scans were acquired to enable the signal to reach steady-state, followed by 136 volumes.

For 13 participants, the manufacturer’s EPI internal reference option was used. The internal reference acquires four non-phase-encoded echoes before the EPI echo train, which are used to apply a phase correction to the EPI data. Upon further inspection of the data this was shown to contribute to slice misalignment in the anterior-posterior direction in 10 of the 13 participants (20 out of 52 task runs) and thus the setting was disabled for subsequently recruited participants. To keep the two MRI visits identical, however, the internal reference was used on both MRI visits for the first 13 participants, even after the issue was discovered.

### 2.4 Data analysis

#### 2.4.1 Behavioural data

Data were analysed using SPSS v29.0.2.0. The outcome measure was maximum hand-grip force sampled during each trial, expressed as a percentage of individual grip strength threshold. Average task performance was assessed across all squeeze blocks. A 2 × 2 repeated measures ANOVA with visit and run as factors compared grip force exerted.

#### 2.4.2 Imaging data

##### 2.4.2.1 Preprocessing

Data were processed using Spinal Cord Toolbox (SCT) versions 6.5 and 7.0 (De Leener et al., 2017), AFNI’s *3dWarpDrive* (Cox, 1996; Cox & Hyde, 1997), and FSL version 6.0.7.14 (Jenkinson et al., 2012; Smith et al., 2004). In particular, SCT version 7.0 was used only for registration steps to facilitate implementation of the newly- introduced rootlet-based registration (Bédard, Valošek, et al., 2025). Visual quality assurance was performed on raw data and at each stage of processing. Twenty task runs acquired with an early version of the functional sequence using the internal reference (see above) had several slices come out of alignment with the rostrocaudal axis of the spinal cord due to a shift in the anterior-posterior (EPI phase-encoding) direction. A custom in-house Matlab version 9.5.0 (Mathworks Inc.) script was used to move the slices back into alignment with the rest of the cord. Briefly, for each slice, a 1D projection along the anterior/posterior direction was calculated for each time-point by summing the voxels in the left/right direction across the spinal cord. The anterior/posterior shift was determined by calculating the maximum of the cross correlation of the projection at each time-point with the first time-point. The shift was then applied to the image data in a block circular manner. Only shifts by an integer number of voxels were applied to avoid the need for an extra interpolation step. This step was performed prior to any other preprocessing.

For all functional data, brainstem structures were separated from cervical volumes at the level of the odontoid process. Subsequently, spinal cord functional data were slice-wise motion-corrected for x and y translations, using an in-house implementation of AFNI’s *3dWarpDrive* following the steps in the Neptune Toolbox (https://github.com/NeptuneToolbox). Motion-corrected data were smoothed with an in-plane 2D Gaussian kernel with full width at half maximum (FWHM) of 2 mm using a custom in-house script relying on tools from AFNI and FSL.

Warping parameters for spatial normalisation to the PAM50 spinal cord template (De Leener et al., 2018) were determined using rootlet-based registration (Bédard, Valošek, et al., 2025). First, *sct_deepseg* was used to segment the cord from the cerebrospinal fluid (CSF); an EPI-specific algorithm was used on the temporal mean of motion-corrected functional data (Banerjee et al., 2025) and a contrast-agnostic algorithm on the T2-weighted structural image (Bédard, Karthik, et al., 2025). Dorsal cervical rootlets were identified on T2-weighted data with *sct_deepseg rootlets* (Valošek et al., 2024). Manual adjustment of cord and rootlet segmentation was performed in FSLeyes as required (McCarthy, 2022). Warping parameters for registration of functional data to the PAM50 template were created by combining warp parameters from: (1) registering the subject-specific structural T2-weighted image and functional data, using manually created disc labels on both images; and (2) registering the T2-weighted image to the PAM50 T2-weighted template via *sct_register_to_template* with rootlet labels (Bédard, Valošek, et al., 2025). Final warping fields were obtained by applying concatenated warps from the previous step to bring the temporal mean of functional data to the PAM50 T2*-weighted image via *sct_register_multimodal* (De Leener et al., 2018). The resultant warping fields were used for PAM50 registration of contrast of parameter estimates (COPE) generated from subject-level modelling with *sct_apply_transfo*.

The Physiological Noise Modelling (PNM) toolbox (Brooks et al., 2008) was used to generate 33 slice-specific regressors accounting for physiological noise based on cardiac and respiratory traces, and CSF signal. CSF masks were created by dilating the spinal cord mask to encompass the surrounding subarachnoid cavity and subsequently subtracting the former from the latter. A boundary of one voxel was maintained between the cord and CSF masks to avoid partial volume effects.

##### 2.4.2.2 Subject-level modelling

Spinal cord activation was assessed using mass univariate general linear models (GLMs) in FSL FEAT, using FILM pre-whitening and highpass temporal filtering at 90 s. GLMs included explanatory regressors for squeeze trials, along with slice-wise nuisance regressors accounting for physiological noise (33 PNM regressors) and in-scan motion (x and y translations). Additionally, volume censoring regressors were added to exclude the first trial of each squeeze block due to large behavioural variability in the responses at the start of the squeeze blocks (see Supplementary Figure 1). Two contrasts identified: (1) motor activity greater than rest (COPE multiplier = 1), and (2) for additional quality assurance and inspection of potential task negative activity – rest greater than motor activity (COPE multiplier = -1).

We also conducted a parametric modulation analysis to account for variability in task performance. These GLMs included a parametric regressor representing the grip force exerted during each squeeze trial, in addition to the regressor modelling squeeze trials and slice-wise nuisance regressors described above. Three contrasts were estimated: (1) motor activity greater than rest, accounting for grip force variability (hereafter referred to as grip force-adjusted motor activity), (2) activity positively correlated with grip force, and (3) activity negatively correlated with grip force.

##### 2.4.2.3 Group-level modelling

Task effects were assessed using a one-sample t-test comprising averages of subject-level COPEs across all four runs of the task (two runs from each visit). Additional group-level maps were computed to assess activation in each run and visit separately. These analyses were conducted for both standard and parametric modulation models.

Group-level analyses were conducted within *a priori* defined ROIs and across the whole cervical cord. ROIs included C5-T1 spinal segmental levels (Frostell et al., 2016), derived from the PAM50 spinal levels atlas and corresponding to the dermatomal and myotomal projection of the hand (Sonoo, 2023; Weber et al., 2020).

##### 2.4.2.4 Impact of number of task runs on group-level activation

To investigate the impact of data quantity on group-level activation maps, four group-level models were conducted. The group-level models included: only one run (visit 1, run 2 was used since no clusters survived FWE correction in visit 1, run 1), two runs (average of runs 1 and 2 from visit 1), three runs (average of runs 1 and 2 from visit 1, and run 1 from visit 2), and all fours runs (average of runs 1 and 2 from visits 1 and 2). These analyses were conducted for standard and parametric modulation models.

##### 2.4.2.5 Statistical inference

All group-level models used non-parametric permutation-based testing implemented in *randomise* (FSL) with threshold-free cluster enhancement (5000 permutations, *p* < 0.05) and family-wise error (FWE) rate correction.

#### 2.4.3 Test-retest reliability

Test-retest reliability was assessed both within and between visits. For within-visit reliability, we compared the mean of the first task run across visits 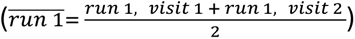 with the mean of the second task run across visits 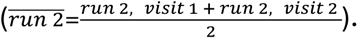 For between-visit reliability, we compared the mean of both task runs from visit 1 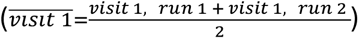 with the mean of both task runs from visit 2 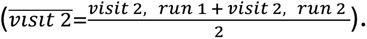

##### 2.4.3.1 Intraclass correlation coefficient

ICC was used to quantify the reliability of both behavioural and imaging data. Following previous recommendations, ICC values were categorised accordingly: <0.4 as *poor*, 0.4–0.59 as *fair*, 0.6–0.74 as *good*, and >0.75 as *excellent* (Cicchetti, 1994; Fleiss et al., 2013).

###### 2.4.3.1.1 Behavioural data

Reliability of task performance was assessed using ICC(3, 2) and calculated in SPSS v29.0.2.0.

###### 2.4.3.1.2 Imaging data

ICC values were calculated for each voxel (i.e. voxelwise) using the locally-developed ICC toolbox (Caceres et al., 2009) running in Matlab version 9.5.0 (Mathworks Inc.). Reliability was calculated for the whole cervical cord (PAM50 cord mask), the activation network, and within spinal segmental levels C5-T1 (PAM50 spinal segmental masks). The activation network was obtained using a one-sample *t*-test of the first measurement (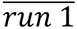 for within-visit comparison, 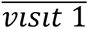 for between-visit comparison) with a voxelwise *t*-statistic threshold of 3.5 (equivalent to *p* = 0.001) conducted in SPM8, as per the original manuscript describing this approach (Caceres et al., 2009). Median ICC values are reported, defined as the reliability measure obtained from the median of the ICC distributions within regions.

In addition to the pre-registered voxelwise approach, test-retest reliability of the average motor contrast estimate was computed. Mean values were extracted from motor-related COPEs from subject-level models from the whole cervical cord (PAM50), the activation network obtained from group-level *randomise* (*p*_FWE_ < 0.05) models (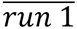for within-visit comparison, 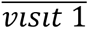 for between-visit comparison), and within spinal segmental levels C5-T1 (PAM50). Reliability of motor activation was assessed in both standard and parametric modulation analysis.

To supplement the assessments of reliability of motor activation and to test the reliability of overall fMRI signal, within- and between-visit ICC was calculated for temporal signal-to-noise ratio (tSNR). tSNR was calculated on motion-corrected data prior to any further preprocessing or modelling steps to avoid artificially inflating the measure. The mean motion-corrected functional image was divided by its standard deviation. Mean tSNR was extracted in native space for the whole cervical cord using subject-specific cord masks generated during preprocessing (see Section 2.4.2.1).

Reliability of average signal and tSNR was assessed using ICC(3, 2) in SPSS v29.0.2.0.

###### 2.4.3.1.3 Dice similarity coefficient

The spatial consistency of group-level activation maps (*p*_FWE_ < 0.05) within and between visits was assessed using Dice similarity coefficient (DSC; Dice, 1945). DSC was calculated using *3ddot* from AFNI 25.2.18 for both motor activation and grip force-adjusted motor activation. DSC ranges from 0 to 1, with higher values indicating better overlap between sets.

## 3 Results

### 3.1 Motor performance

Participants produced consistent hand-grip force during the squeeze blocks (mean across all task runs ± SD = 83.50% ± 12.40; visit 1, run 1 = 84.64% ± 16.89; visit 1, run 2 = 79.45% ± 13.62; visit 2, run 1 = 88.43% ± 14.47; visit 2, run 2 = 81.67% ± 12.88). No grip force was detected during rest blocks (Figure 2).

**Figure 2.**
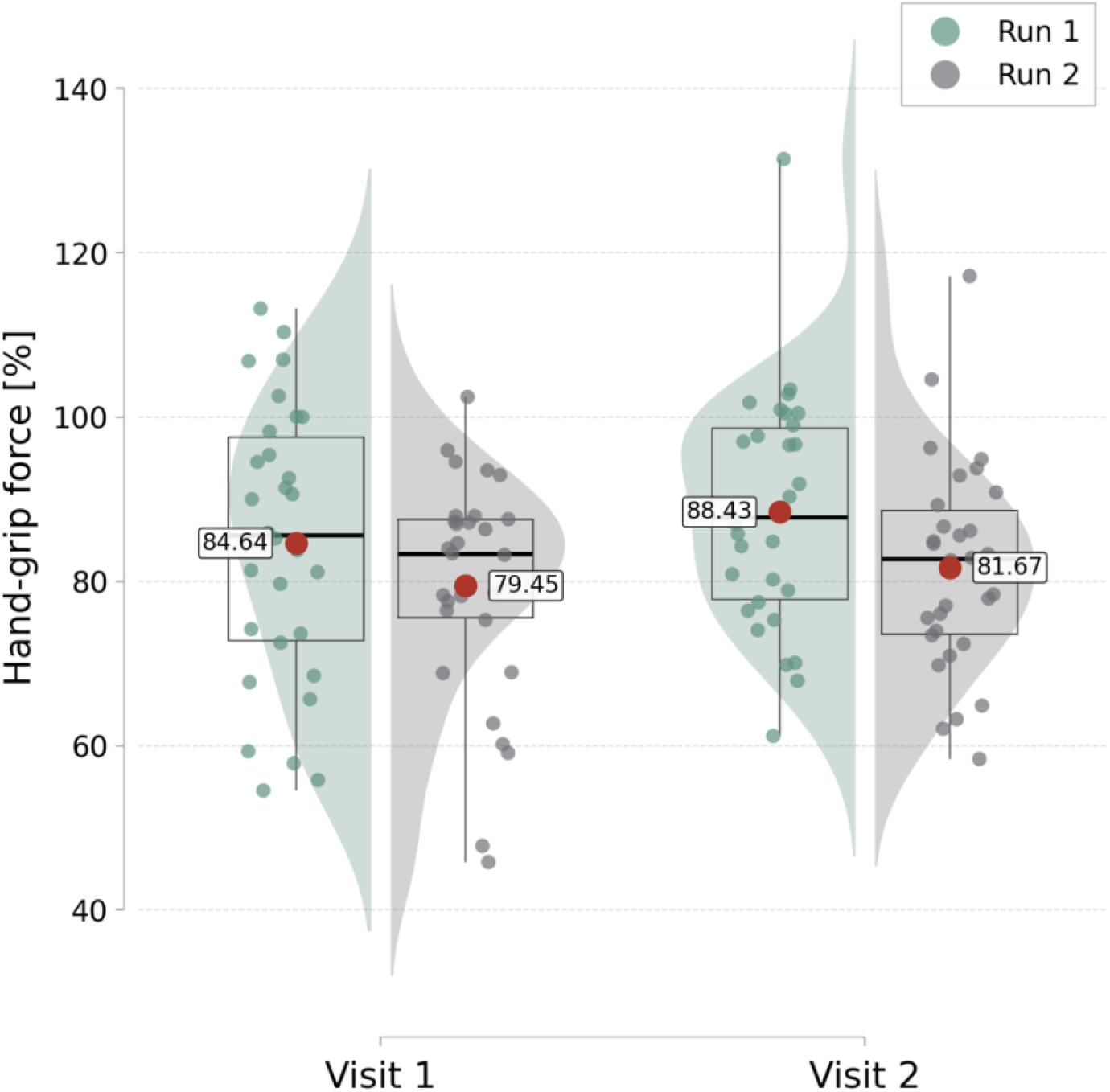
Average hand-grip force exerted by participants during each of the task runs on each visit. The values are expressed as percentage of individually thresholded grip force. Boxplots mark the median (thick black line), interquartile range (box), and minimum and maximum values (whiskers). Red dots and the associated labels mark the mean.

No significant differences were observed between visits (F(1, 29) = 3.14, *p* = 0.087, η²p = 0.098), however, lower grip force was observed in the second run of the task across both visits (F(1, 29) = 8.58, *p* = 0.007, η²p = 0.228). No interaction effects between visit and run were found (F(1, 29) = 0.23, *p* = 0.631, η²p = 0.008).

### 3.2 Motor activation

Group averaging across all four task runs (two runs per visit, two visits) showed ipsilateral right-lateralised motor-related activation in both ventral and dorsal regions of segmental levels C5-C8, as well as bilateral ventro-dorsal activation at levels C2 and C3 when assessed across the whole cervical spinal cord (Figure 3A). A similar extent of ipsilateral ventro-dorsal activation was observed when using *a priori* defined ROIs as masks at spinal segmental levels C5, C6, C7, and T1 (Figure 3B). Analysis of grip force-adjusted motor activation yielded similar results, with additional bilateral activation observed at C7 and right activation at T1 in whole cord and ROI analyses, and right ventro-dorsal activation in ROI analysis of level C8 (Supplementary Figure 2).

**Figure 3.**
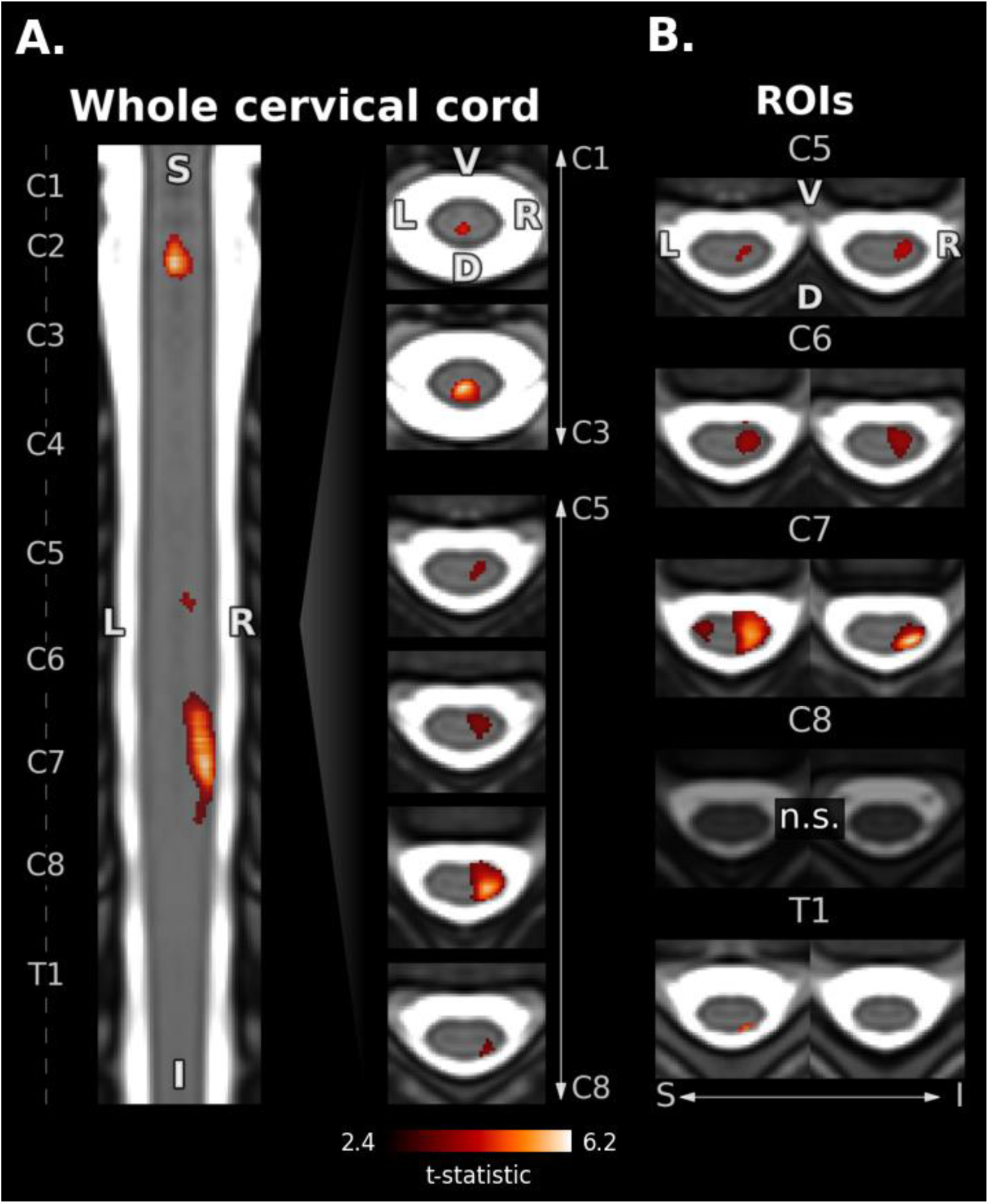
Group-level motor activation averaged across all task runs. Statistically significant t-statistics are shown (*p*_FWE_ < 0.05). Labels denote spinal segmental levels. **A.** Activation obtained from whole cervical cord analysis shown on one representative coronal slice and representative axial slices from spinal segmental levels C2-C3 and C5-C8. **B.** Activation obtained from ROI analysis at each spinal segmental level C5-T1. D = dorsal, I = inferior, L = left, n.s. = non-significant, R = right, ROI = region of interest, S = superior, V = ventral.

When assessing group-level activation for each task run separately within the whole cord mask, motor-evoked activation was observed in ipsilateral ventro-dorsal regions spanning levels C6-C7 (run 2, visit 1; runs 1 and 2, visit 2; Figure 4). ROI analyses of individual task runs, showed primarily ipsilateral ventro-dorsal activation at level C7 (run 2, visit 1; runs 1 and 2, visit 2), with some clusters spanning bilaterally. Additional activation clusters were observed bilaterally at C6 (run 1, visit 1) and ipsilaterally at C5 (run 2, visit 2) and C8 (run 1, visit 2; Figure 5). Grip force-adjusted motor activation was observed in the same regions, with an additional right ventro-dorsal activation at level C8 (run 1, visit 1) and medial activation at C5 (run 2, visit 2) in whole cord analysis (Supplementary Figure 3). In addition to the clusters observed in the main analysis, grip force-adjusted ROI analysis revealed primarily ventral ipsilateral clusters at C6-C8 in run 1 of visit 1 and medial activation at C6 in run 2 of visit 2 (Supplementary Figure 4).

**Figure 4.**
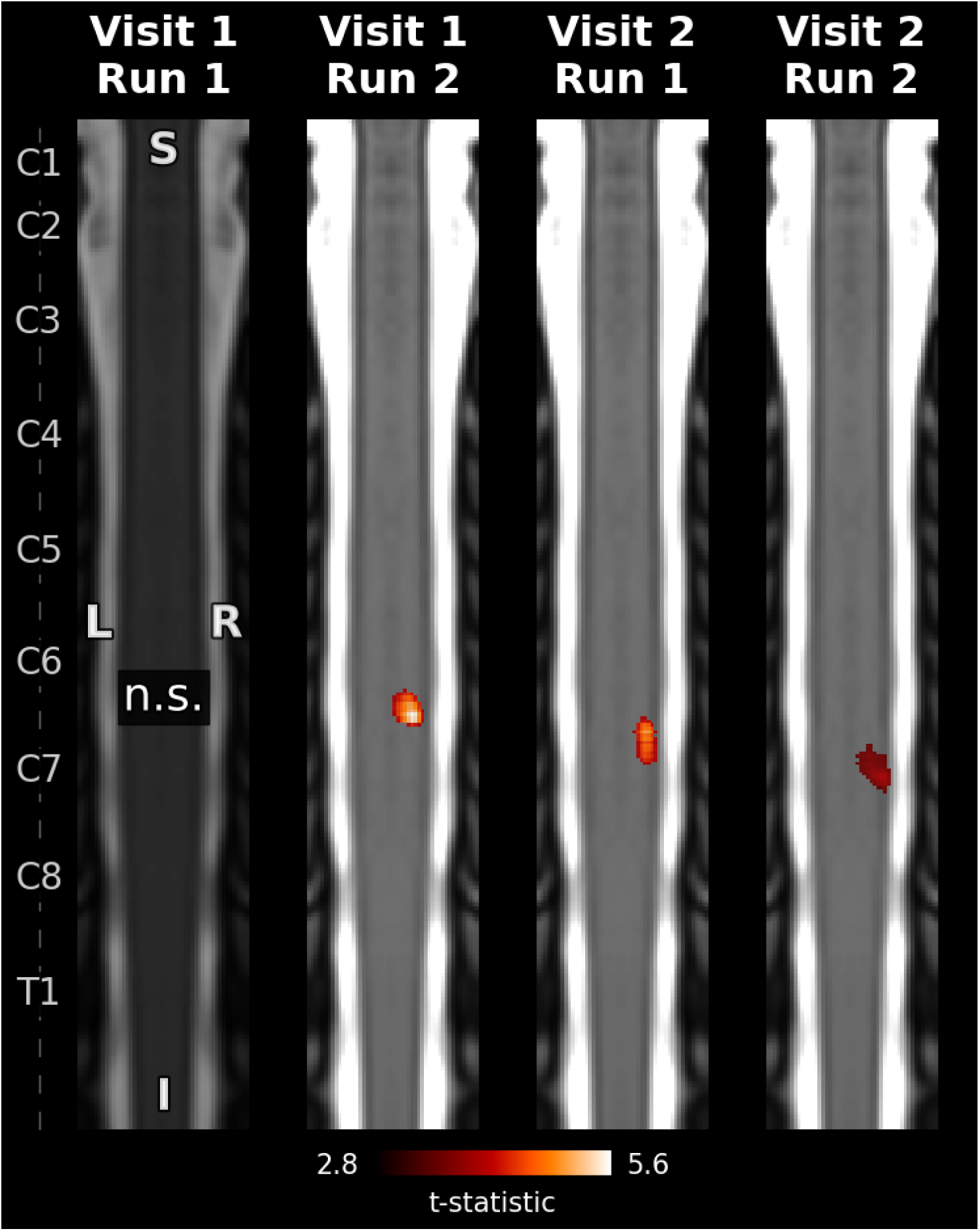
Group-level motor activation in each run of the task assessed within the whole cervical cord. One representative coronal slice per task run is presented. Statistically significant t-statistics are shown (*p*_FWE_ < 0.05). Left axis denotes spinal segmental levels. I = inferior, L = left, n.s. = non-significant, R = right, S = superior.

**Figure 5.**
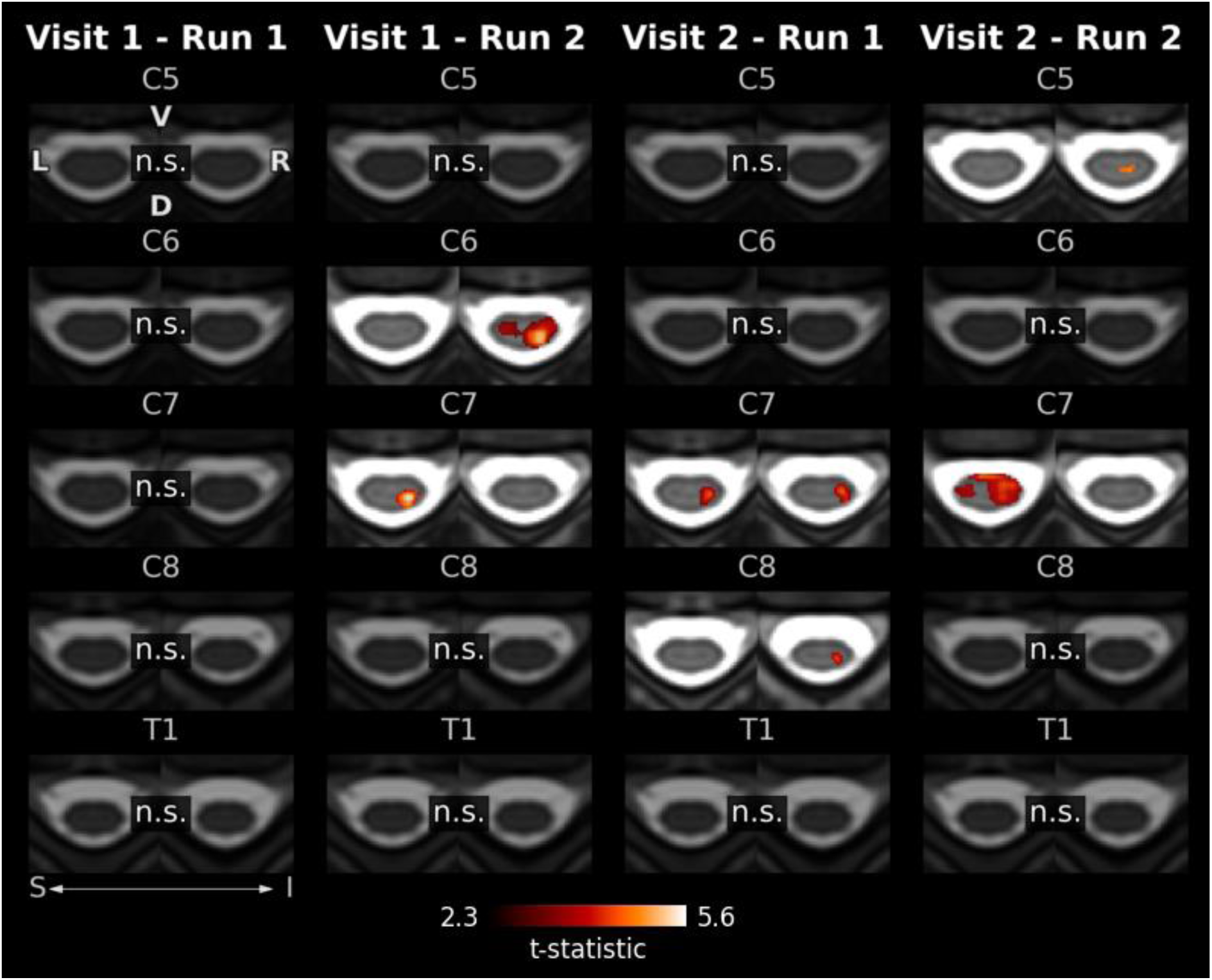
Group-level motor activation in each run of the task assessed within each spinal segmental level ROI (C5-T1). Statistically significant t-statistics are shown (*p*_FWE_ < 0.05). D = dorsal, I = inferior, L = left, n.s. = non-significant, R = right, ROI = region of interest, S = superior, V = ventral.

No significant clusters were detected in the contrast exploring activation during rest periods, or when exploring the relationship between grip force and cord activation.

### 3.3 Impact of data quantity on motor activation

To investigate the impact of data quantity on motor activation, we repeated group-level models, varying the number of task runs included from one to four. We conducted this comparison on both motor and grip force-adjusted motor activation. In both modelling approaches, increasing the number of task runs included in each analysis resulted in more active voxels (*p*_FWE_ < 0.05) across the cord (motor activation, r = 0.96, *p* = 0.042, 95% CI [-0.04; 1.00]; grip force-adjusted motor activation, r = 0.95, *p* = 0.046, 95% CI [-0.09; 1.0]) and skewed the distribution of statistically significant t-values towards lower estimates (Figure 6B and Supplementary Figure 5B). Increasing the number of runs also led to a broader spatial extent of activated voxels across modelling approaches, including at expected segmental levels C6-C8 (Figure 6A and Supplementary Figure 5A).

**Figure 6.**
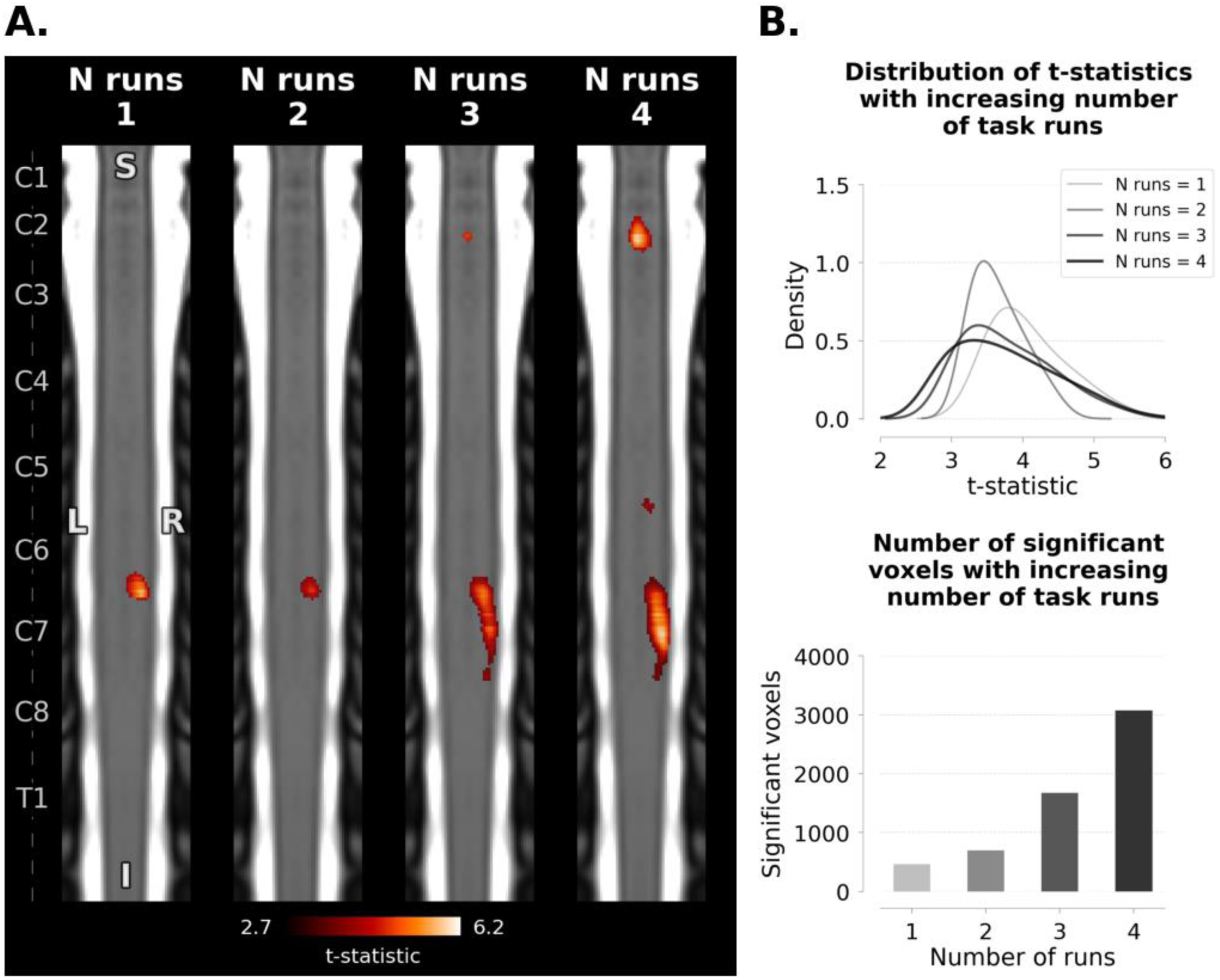
Investigation of the impact of increasing the number of task runs included in analysis on group-level motor activation across the cervical spinal cord (*p*_FWE_ < 0.05). **A.** Group-level motor activation maps. One representative coronal slice per group-level is presented. Left axis denotes spinal segmental levels. **B.** Graphs showing the density of t-statistics (top) and number of active voxels (bottom) across the cord with increasing the number of task runs. I = inferior, L = left, N = number, R = right, S = superior.

### 3.4 Test-retest reliability

#### 3.4.1 Intraclass correlation coefficient

##### 3.4.1.1 Behavioural data

Reliability of task performance was *good* within visit (ICC = 0.73) and *excellent* between visits (ICC = 0.84).

##### 3.4.1.2 Imaging data

Voxelwise reliability estimates of motor activation were *poor*-to-*fair* for both within- and between-visit comparisons (ICC range = 0.08-0.44; Figure 7). Within- and between-visit reliability of average regional activation was *poor*-to-*fair* in most ROIs (ICC range = 0-0.59), with only a few exceptions. *Good* within-visit reliability of COPE average was observed at C5 (ICC = 0.62) and C6 (ICC = 0.62), as well as between-visit across the cervical cord (ICC = 0.69).

**Figure 7.**
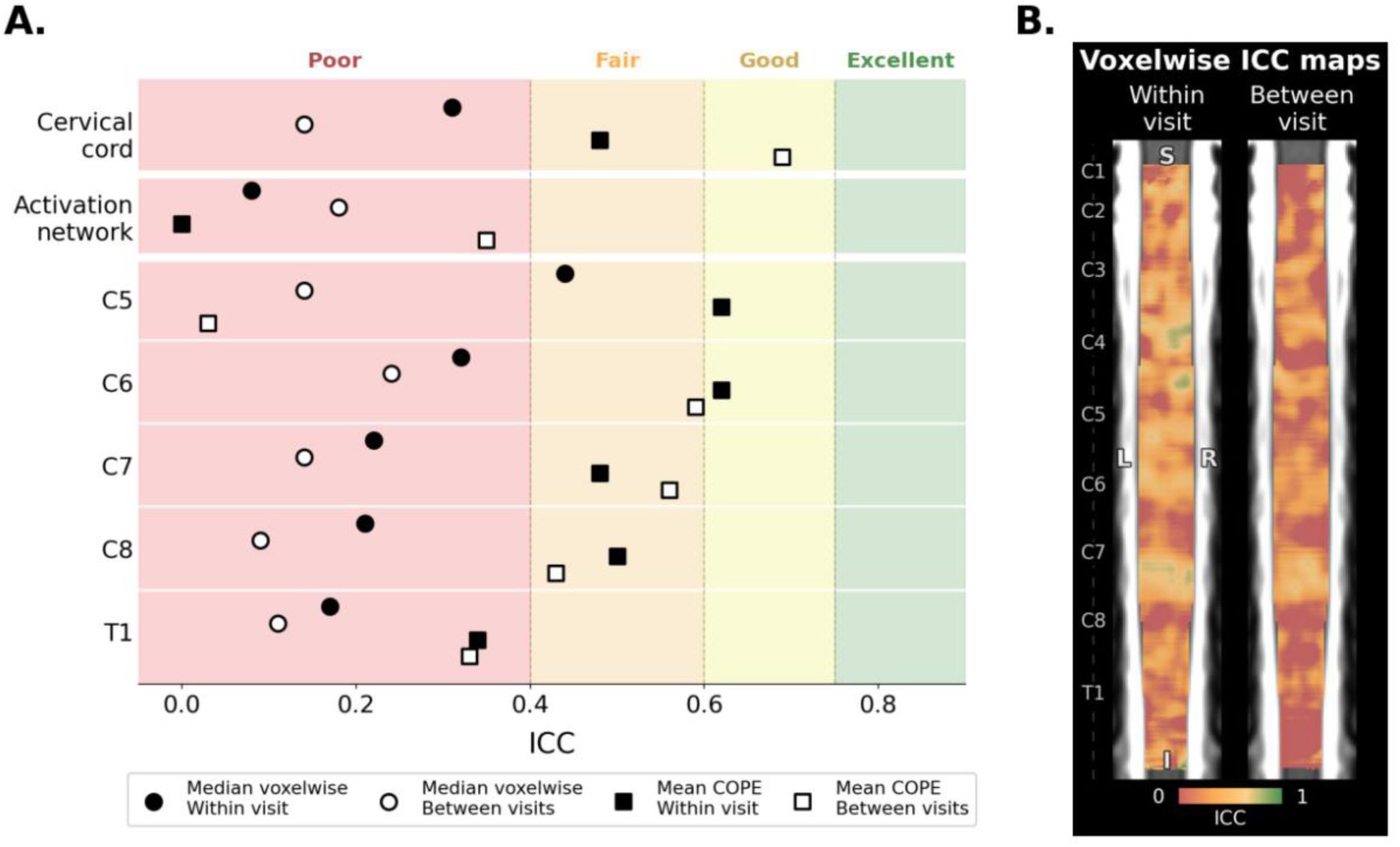
ICC estimates for within- and between-visit reliability of motor activation. **A.** Forest plot showing median ICC for voxelwise assessments and ICC of mean COPE across the cervical spinal cord, the motor activation network, and spinal segmental levels C5-T1. **B.** Statistical maps showing ICC distribution for individual voxels within the cervical cord for within- and between-visit reliability assessments. COPE = contrast of parameter estimates, ICC = intraclass correlation coefficient, SE = standard error.

Accounting for grip force (grip force-adjusted motor activation) lowered ICC estimates by an average of 0.03 (t(27) = 2.4, *p* = 0.025, Cohen’s d = 0.45, 95% CI [0.05; 0.83]; Supplementary Figure 2).

In addition to motor activation, we also assessed the reliability of tSNR as a measure of overall signal acquired during scanning. Average tSNR across all task runs was 27 ± 6.01 (visit 1, run 1 = 26.43 ± 6.63; visit 1, run 2 = 26.62 ± 6.51; visit 2, run 1 = 27.46 ± 5.16; visit 2, run 2 = 27.5 ± 5.86). tSNR showed *excellent* reliability both within- (ICC = 0.96) and between-visit (ICC = 0.79).

##### 3.4.1.3 Dice similarity coefficient

DSC assessed the spatial overlap of group-level activation within- and between-visit, indicating poor overlap for both motor activation (DSC within-visit = 0.01, DSC between-visit = 0.04) and grip force-adjusted motor activation (DSC within-visit = 0.14, DSC between-visit = 0.12).

## 4 Discussion

This study investigated cervical spinal cord activation during a unilateral hand grasping motor task and its test-retest reliability. We observed the expected ipsilateral activation in ventro-dorsal regions spanning spinal segmental levels C5-T1, with additional bilateral ventro-dorsal and medial activation at segments C2-C3. Despite *good*-to-*excellent* reliability of task performance and overall fMRI signal (tSNR), reliability of motor activation was primarily within the *poor*-to-*fair* range for voxelwise and mean signal estimates, with little to no spatial overlap in activation maps across task runs. Herein we discuss these findings and consider the challenges in establishing reliability of measurements, emphasising the pressing need to distinguish between true measurement error and biologically meaningful sources of variability within individuals.

### 4.1 Hand grasping evoked ipsilateral ventro-dorsal activation in spinal segmental level C6-C8

The first aim of this study was to characterise motor-evoked cervical spinal cord activation. The ventro-dorsal activation observed at spinal segmental levels C5-T1 was largely lateralised to the hand executing the movement, with some clusters extending medially. This ipsilateral predominance aligned with our predictions and the existing literature. Spinal cord anatomy is inherently lateralised, with afferent and efferent projections ipsilateral to the corresponding body part (Ganapathy et al., 2025). Previous spinal fMRI studies using unilateral motor tasks have reported mostly ipsilateral activation, which aligns with established spinal circuitry (Barry et al., 2021; Braaß et al., 2023; Hemmerling et al., 2023; Maieron et al., 2007; Weber et al., 2016b). However, additional bilateral and/or medial activation has also been observed during unilateral movements (Bouwman et al., 2008; Hemmerling et al., 2023; Oliva et al., 2025; Stroman & Ryner, 2001; Vahdat et al., 2015), similar to that observed in the medial parts of levels C5-C7 in our ROI analysis. This medial activation likely reflects neural processing in the intermediate zone, a gray matter region between the ventral and dorsal horns that connects the left and right hemicords and houses commissural interneurons (Ganapathy et al., 2025). Electrophysiology recordings in non-human primates have shown that commissural interneurons are involved in hand grasping, facilitating motor coordination of the upper limb (Soteropoulos et al., 2013; Takei & Seki, 2010). Reports from human studies show that analogous processes in the lumbar cord may underpin motor coordination during lower limb movements (Hanna-Boutros et al., 2014; Mrachacz-Kersting et al., 2018). Nonetheless, it is important to note that methodological choices, such as in-plane voxel size and associated partial volume effects, along with spatial smoothing, limit the spatial specificity of spinal fMRI. Consequently, while medial activation may reflect commissural interneuron activity, finer spatial resolution is required to validate this interpretation.

In line with our hypothesis, motor-evoked activation observed in this study encompassed both ventral and dorsal regions of the cord. The ventral and dorsal gray matter horns primarily comprise motoneurons and sensory neurons, respectively (Ganapathy et al., 2025). Both ventral and dorsal activation are consistently reported in spinal fMRI studies probing motor function, with peaks often located more ventrally (Landelle et al., 2021).

Recruitment of both ventral and dorsal regions of the cord is expected, given the nature of the sensorimotor task used here, the complex circuitry of the cord, and the intricate connections among motoneurons, sensory neurons, and interneurons, which give rise to various reflex loops and facilitation-inhibition interdependencies (Pierrot-Deseilligny & Burke, 2012). Co-activation of ventral and dorsal horns might thus reflect sensory feedback accompanying movement or early sensorimotor integration (Enoka, 2008; Landelle et al., 2021; Yeganegi et al., 2018). Despite a somewhat larger voxel size than typically used in the field (1.41 × 1.41 mm versus 1 × 1 mm) and the use of spatial smoothing lowering the spatial specificity of our findings, studies using higher spatial resolution (Hemmerling et al., 2023; Oliva et al., 2025; Weber et al., 2016b) and not incorporating spatial smoothing (Hemmerling et al., 2023) reported similar activation patterns, supporting the interpretation that hand grasping recruits diffuse areas within spinal cord segments.

In agreement with established muscle innervation pathways (Schirmer et al., 2011), hand grasping led to activation at spinal segmental levels spanning C5-T1, with peaks at levels C6 and C7, consistent with our hypothesis. Hand grasping involves muscle groups innervated by median and ulnar nerves of the brachial plexus, which receive projections from C5-C8 and C8-T1 nerve roots, respectively (Dawson-Amoah & Varacallo, 2025; Schirmer et al., 2011). Previous spinal fMRI studies involving similar tasks have shown analogous activation spanning C5-T1 (Backes et al., 2001; Barry et al., 2021; Braaß et al., 2023; Giulietti et al., 2008; Hemmerling et al., 2023; Islam et al., 2019; Kinany et al., 2019; Ng et al., 2006; Oliva et al., 2025; Stroman et al., 1999; Stroman & Ryner, 2001; Weber et al., 2016b), with peaks for wrist movements occurring at C6-C7 and finger movements at C6-T1 (Landelle et al., 2021). The activation we observed at levels C5-T1 when combining all task runs, aligns with the existing literature and the known somatotopic organisation of spinal cord function.

### 4.2 Hand grasping involved additional activation in rostral cervical cord segments

In addition to the expected recruitment of spinal levels C5-T1, we observed activation at spinal segmental levels C2-C3. Although previous studies have documented activation spreading to adjacent segments, few have reported distinct clusters in such rostral regions (Landelle et al., 2021). Engagement of C2-C4 spinal segmental levels has been observed in addition to the expected dermatomal/myotomal projections during finger tapping (Govers et al., 2007; Ng et al., 2008) and C3-C4 during sensory stimulation of the fingers (Stracke et al., 2005). Drawing from electrophysiology literature, activation of rostral segments of the cervical spine may be underpinned by the involvement of neuronal projections spanning multiple spinal segmental levels, which are involved in somatosensory integration, proprioception, and fine motor control (Flynn et al., 2011; Pierrot-Deseilligny & Burke, 2012). This interpretation is further supported by the location of the peaks of our C2-C3 activation mapping to the intermediate zone, which contains interneurons and propriospinal neurons (Ganapathy et al., 2025). Additionally, C2-C4 segments receive projections from the posterior head, as well as the neck and shoulders (Schirmer et al., 2011; Walji & Tsui, 2016). We speculate that the activation observed at these levels may reflect additional sensory feedback and/or contraction of muscles not directly involved in hand grasping, but that aid in maintaining appropriate posture during scanning.

Relating this finding to the spinal fMRI literature is complicated by the fact that the studies reporting rostral activation have been conducted with modest sample sizes (N = 9-12) and prior to the many developments in optimising spinal fMRI data acquisition and processing that have occurred in the last decade (Kinany et al., 2022; Powers et al., 2018). The consistency of these findings is difficult to establish given that the majority of current spinal fMRI literature focuses on spinal segmental levels C4-T1, corresponding to the afferents and efferents of the upper limb, either by using a smaller field of view (Barry et al., 2021; Hemmerling et al., 2023; Oliva et al., 2025; Weber et al., 2016b) or by restricting analyses to this area (Kinany et al., 2019; Vahdat et al., 2015). Despite these considerations, and given the specificity of the activation to motor periods of the task, the C2-C3 activation observed here likely reflects neurophysiological processes related to movement.

### 4.3 Poor test-retest reliability of motor-activation can stem from methodological choices

Our second objective was to assess the test-retest reliability of spinal motor-evoked activation. Despite highly reliable task performance and tSNR (ICC > 0.7), motor activation reliability was substantially lower than expected (average ICC = 0.3), falling primarily within the *poor*-to-*fair* range. Similarly, the spatial representation of activation maps, particularly their rostrocaudal extent, varied across task runs. Furthermore, despite reliability estimates typically being higher for assessments conducted within the same visit, as compared to those conducted during different visits, this pattern was not consistent in our data, except for tSNR (Bennett & Miller, 2010, 2013).

Our reliability findings align with previous spinal fMRI studies reporting *poor* test-retest reliability in motor (Weber et al., 2016b), sensory (Dabbagh et al., 2024; Weber et al., 2016a), and resting-state paradigms (Kowalczyk et al., 2024). These findings parallel the broader neuroimaging literature, where brain fMRI studies similarly report modest reliability (average ICC = 0.5) (Bennett & Miller, 2010; Elliott et al., 2020; Kragel et al., 2021; Noble et al., 2017, 2019) and tSNR is notably higher (typically three times that found in the cord; Murphy et al., 2007). Measurement error is a large contributor to low reliability, and considering that spinal fMRI faces additional technical challenges leading to greater signal variability (Kinany et al., 2022; Powers et al., 2018), perhaps low ICC values should be expected. The small diameter of the cord, impact of physiological noise, close proximity of different tissue types, and additional in-scan motion related to CSF pulsation can all reduce the signal-to-noise ratio of fMRI data obtained from the spinal cord (Kinany et al., 2022; Landelle et al., 2021; Powers et al., 2018). Indeed, the impact of noise on spinal fMRI reliability has been extensively characterised recently, underscoring the importance of accounting for the various sources of noise present in spinal fMRI recordings (Kaptan et al., 2022).

Nonetheless, low reliability does not equate to low validity. Past work has shown little association between reliability and the ability to predict behaviour or reidentify individuals from intrinsic brain activity, demonstrating that data with low reliability still contains biologically meaningful information (Noble et al., 2017). Furthermore, preprocessing approaches developed to account for artefactual signal and thus boost validity, including in-scan motion or unrelated physiological processes, have been shown to reduce test-retest estimates in both brain and spinal fMRI (Birn et al., 2014; Kaptan et al., 2022; Lipp et al., 2014; Noble et al., 2017, 2019). This likely reflects more structured properties of noise (e.g. cardiac cycle, respiration, CSF pulsation) compared to the more dynamic nature of neural processing (Kinany et al., 2020). Consequently, while continued efforts to improve image acquisition, preprocessing, and modelling approaches will undoubtedly increase both the reliability and validity of spinal fMRI (Barry et al., 2016; Kaptan et al., 2022), it is also important to consider that observations of low reliability may be underpinned by biologically meaningful variability in the measured signal.

In simple terms, ICC is the ratio of between-participant to within-participant variance (Fleiss et al., 2013). Consequently, a high ICC is obtained when the variance between participants is large, while variance within participants is minimal. Traditionally, however, fMRI tasks are designed to minimise between-participant variance and elicit a robust group-level response (Fröhner et al., 2019; Hedge et al., 2018). Even highly repeatable group-level activation maps are not always underpinned by reliable individual-level activation maps (Fröhner et al., 2019). While the lower level of measurement error at group level, compared to individual level, might play a part in this phenomenon, simulation and longitudinal analyses of brain fMRI suggest an important role of trait- and state-related individual differences (Fröhner et al., 2019; Hedge et al., 2018). The brain and spinal cord form a dynamic system, constantly adapting to external and internal demands (Finn & Rosenberg, 2021; Kaptan et al., 2024; Kinany et al., 2020; Uddin, 2020). Low ICC values and variable activation patterns across task runs might be explained by fluctuating psychological and neurophysiological states within individuals. To date, validated mixed-effects approaches, which consider within-, as well as between-individual variability, remain reliant on random field theory assumptions for statistical inference (Worsley, 2003). Important assumptions of stationary smoothness across the image volume are violated by the inherently anisometric shape of the cord, rendering their application invalid for spinal fMRI. Nonetheless, biologically meaningful variability influencing our data is in line with *good*-to-*excellent* reliability of tSNR observed here, indicating that while the spatial extent and magnitude of responses related to hand grasping was variable, the overall signal remained stable over time.

### 4.4 Variability in motor activation has likely behavioural and neurophysiological bases

#### 4.4.1 Behavioural variability

It is possible that the variability of activation patterns and magnitude reflects differences in performance across task runs, such as the exerted force or recruited muscle groups. Motor-related intensity coding has previously been observed in the spinal cord using fMRI, showing that activation scales positively with the force exerted (Braaß et al., 2023; Madi et al., 2001; Oliva et al., 2025). While all participants in our study achieved adequate performance in all task runs (approximately 80% of individualised grip force), the grip force they exerted in the second run of the task across both visits was approximately 6% lower than that exerted in the first runs. This suggests a potential effect of fatigue on task performance, inherent to conducting multiple identical assessments within the same testing session and potentially leading to the lower test-retest reliability of performance within- compared to between-visits. Nonetheless, the similar pattern and magnitude of motor activation across task runs observed in the analysis accounting for grip force (grip force-adjusted motor activation), suggests that trial-by-trial variability in evoked grip force alone is an unlikely explanation for the differences in activation patterns across task runs.

Given that hand grasping involves contractions of multiple muscle groups, including those of the hand and forearm (Dawson-Amoah & Varacallo, 2025; Schirmer et al., 2011), the variability in activation may represent differences in motor control strategies employed by participants across task runs. Combining spinal fMRI with electromyography has demonstrated distinct segmental localisation of motor-evoked activity depending on the type of movement performed, such as wrist extension (C5-C7), wrist adduction (C7-C8), and finger abduction (C7-T1) (Kinany et al., 2019). Since multiple strategies can be employed to exert adequate grip-force in the task, incorporating electromyography recordings would have allowed us to track the recruitment of specific muscle groups during the task and identify any potential changes in motor control strategies over time. We recommend the inclusion of orthogonal readouts in imaging studies as an index of behavioural variability, facilitating the capture of associated neural processing with improved precision (Nebe et al., 2023).

#### 4.4.2 Dynamics of neural processing

The variability in activation maps across task runs may also be explained by the plastic and dynamic nature of neural processing in the spinal cord. Motor skill learning has been shown to modulate spinal cord activity (Khatibi et al., 2022; Kinany et al., 2023; Vahdat et al., 2015), and among other things, to induce a caudal-to-rostral shift in activation over time (Khatibi et al., 2022). While previous spinal fMRI literature has focussed on functional plasticity induced by repetition of complex movements, repetition of even simple movements, such as fist clenching, is known to lower activation in the brain over time (Dinstein et al., 2007; Hamilton & Grafton, 2008). Although not explicitly investigated here, we speculate that the differences in activation patterns observed across task runs may be partly attributed to similar repetition suppression taking place at the level of the cord. These insights may explain the inconsistency in the spatial extent of activation across runs. Furthermore, the caudal-to-rostral shift in significant clusters over the course of task runs within each visit observed in our grip force-adjusted analysis echoes that reported following complex motor skill learning, potentially signifying the development of more efficient motor control with practice (Khatibi et al., 2022). Consequently, changing activation patterns observed across task runs may reflect dynamically adapting neurophysiological processes taking place within the spinal cord to exert appropriate motor control. This interpretation challenges the historical view of the spinal cord as a stable information relay and highlights the importance of accounting for the dynamic nature of neural processing in study design.

#### 4.4.3 Individual differences in neuroanatomy

There is known individual variability in spinal cord structure. Spinal cord segmental levels, which describe the rostrocaudal organisation of the spinal cord, are identified by nerve rootlet connection points. By contrast, vertebral levels characterise the rostrocaudal extent of the bony structure of the spine (De Leener et al., 2018; Landelle et al., 2021). Given the ease of vertebral identification and the availability of automated tools, the majority of currently used spatial normalisation pipelines for spinal fMRI rely on vertebral levels (Azad et al., 2021; Bozorgpour et al., 2023; Gros et al., 2019; Rouhier et al., 2020; Ullmann et al., 2014; Vania & Lee, 2021). However, there is a disparity between spinal and vertebral levels, with evidence from postmortem (Lang & Bartram, 1982) and imaging studies (Cadotte et al., 2015) reporting individual differences in nerve rootlet distribution. We suggest that variability in spinal cord architecture is another likely contributor to the inconsistency in the rostrocaudal extent of activation observed in this study and beyond. To minimise the impact between-individual structural differences, we used the recently-introduced rootlet-based template registration (Bédard, Valošek, et al., 2025) with automatic rootlet segmentation (Valošek et al., 2024). Although this method has been shown to improve between-individual alignment and increase the sensitivity of spinal fMRI (Bédard, Valošek, et al., 2025), its impact on spinal fMRI test-retest reliability has yet to be systematically assessed.

### 4.5 Further considerations regarding methodological choices

We have discussed several methodological and biological factors that can affect group-level activation maps and their test-retest reliability. These factors may act in isolation or interact, their effects likely magnified by the unique challenges associated with acquiring fMRI signals from the cord. The complexities of spinal fMRI have been adroitly and exhaustively reviewed elsewhere to which we refer the reader (Kaptan et al., 2024; Kinany et al., 2022; Landelle et al., 2021; Powers et al., 2018). Here, we consider select methodological challenges affecting spinal fMRI research, including statistical power, choice of fMRI task, and optimisation of imaging paradigms.

#### 4.5.1 Number of observations per individual

An important secondary finding of this study concerns the importance of appropriate statistical power in spinal fMRI studies (Murphy et al., 2007). While recent years have seen more than doubling in sample sizes (Braaß et al., 2023; Dabbagh et al., 2024; Hemmerling et al., 2023; Kaptan et al., 2022; Kowalczyk et al., 2024; Landelle et al., 2021; Oliva et al., 2022), power depends not only on the number of participants but also on the amount of observations per participant (Dabbagh et al., 2024; Hemmerling et al., 2023; Weber et al., 2016b). Recent findings from a similar motor paradigm to that used here demonstrated that increasing the amount of observations per participant in spinal fMRI may be more effective at maximising detected activation than simply expanding sample size (Hemmerling et al., 2023), aligning with early reports from motor (Weber et al., 2016b) and sensory studies (Weber et al., 2016a). Such an approach also echoes recommendations reported in brain fMRI, in which a greater number of observations per individual increases both reliability and validity (Noble et al., 2017). Increasing the number of observations can improve accuracy by reducing the prevalence of random unstructured noise originating from transient factors, such as alertness, fatigue, head and neck positioning, thereby distilling signal reflecting stable biological traits and processes (Elliott et al., 2021). Our findings support this approach, showing more robust activation maps when multiple task runs are combined. Furthermore, underpowered imaging studies are prone to overestimating effect sizes (Cremers et al., 2017), a pattern recently confirmed in spinal fMRI (Hemmerling et al., 2023). Extending this evidence, we observed inflated t-statistics when fewer observations per participant were analysed, highlighting that this bias can arise not only from limited sample sizes but also insufficient quantity of data per participant. Consequently, we accord with previous recommendations to acquire more data per individual within adequately powered sample sizes to improve spinal fMRI precision (Hemmerling et al., 2023).

#### 4.5.2 Optimisation of functional paradigms

A well optimised fMRI paradigm must balance technical efficiency in capturing and isolating the BOLD signal and *human* factors, such as attention, habituation, and fatigue. Given the lower effect sizes and greater impact of extraneous noise in the cord compared to the brain, one might approach optimising functional paradigms by maximising signal-to-noise ratio (e.g. by increasing stimulus repetition and/or scan duration; Landelle et al., 2021; Powers et al., 2018; Stroman et al., 2014). This focus on technical optimisation, however, may inadvertently lead to dismissal of large portions of biologically relevant, within-individual variability as noise. fMRI task choice remains a critical design consideration, with particular emphasis on the trade-offs likely in optimising larger versus more specific cord activation. Here, we selected hand grasping due to the large-scale engagement of the upper limb and the large effect sizes associated with the task facilitating assessment of test-retest reliability (Dawson-Amoah & Varacallo, 2025; Landelle et al., 2021; Schirmer et al., 2011). Nonetheless, we reflect that the many strategies employable in executing hand grasping may also contribute to the variability of motor activation observed across task runs, including the spread in rostrocaudal extent. A task choice targeting particular myotomes may offer greater spatial specificity (Kinany et al., 2019) and more consistent cord activation over time.

#### 4.5.3 Outlook

Despite some three decades of development, functional imaging remains in its relative infancy in characterising neurophysiological and psychological sources of within-individual variability. Studies searching for reliable trait-level indices of neural processing have predominated, inadvertently biasing against the largely accepted view of the CNS as a plastic and dynamic system (Finn & Rosenberg, 2021; Kaptan et al., 2024; Kinany et al., 2020; Ricchi et al., 2025; Uddin, 2020). We suggest that a thorough understanding of variability both *between* and *within* complex biological systems is required in spinal fMRI and, more broadly, functional imaging per se. Current paradigms, optimised for group-level inference, offer only limited clinical utility in understanding individual neural processing, for example, those relating to physical trauma, disease progression, or treatment response and recovery. Spinal fMRI still requires ongoing technical improvement and optimisation, but our opinion is that this should not be at the expense of understanding both group- and individual-level signal variability. The combination of traditional fMRI approaches with novel fingerprinting and dense sampling frameworks is a promising avenue for characterising fine-grained spatial and temporal properties of neural function (Finn & Rosenberg, 2021; Michon et al., 2022; Naselaris et al., 2021; Ricchi et al., 2025). We suggest that validating the utility of such approaches in preliminary healthy volunteer studies should catalyse the deployment of spinal fMRI in clinical research.

## 5 Conclusion

Our findings reveal motor-related activation in ipsilateral ventro-dorsal regions of the cervical spinal segments C5-T1, consistent with established dermatomal and myotomal organisation. We also detected activation in segments C2-C3, likely reflecting movement-related sensory feedback, proprioceptive, or postural control mechanisms. Despite consistent task performance and stable overall fMRI signal, the reliability of motor-evoked activation was low. These findings highlight an ongoing challenge in disentangling the impact of measurement error from within-individual variability in understanding reliability in imaging research.

## Data and code availability

The participants of this study did not give written consent for their data to be shared publicly, therefore raw study data are not available.

Spatial maps generated from group-level models and reliability assessments are available on NeuroVault: https://identifiers.org/neurovault.collection:23114. Please note that currently NeuroVault does not support a spinal cord (PAM50) template for online visualisation of spatial maps. Consequently, we recommend downloading the maps and overlaying them on a PAM50 spinal cord template in your software of choice.

All code used in the analysis of this data is openly accessible on the project’s Open Science Framework (OSF) repository: https://osf.io/fjasd/ and GitHub: https://github.com/oliviakowalczyk/spain_squeeze. The data that support the findings of this study are available on request from the corresponding author. The data are not publicly available due to privacy or ethical restrictions.

## Author contributions

Conceptualization: O.S.K., S.M., D.T., S.C.R.W., J.C.W.B., D.J.L., and M.A.H.

Data curation: O.S.K, A.V.

Formal analysis: O.S.K., A.V.

Funding acquisition: S.C.R.W., J.C.W.B., and M.A.H.

Investigation: O.S.K., S.M., and M.A.H.

Methodology: O.S.K., S.M., D.T., J.C.W.B., D.J.L., and M.A.H.

Project administration: O.S.K., S.M., and M.A.H.

Resources: O.S.K., S.M., and M.A.H.

Software: O.S.K. and D.J.L.

Supervision: M.A.H.

Validation: O.S.K., S.M., J.C.W.B., D.J.L., and M.A.H.

Visualization: O.S.K. and A.V.

Writing – original draft: O.S.K.

Writing – review and editing: O.S.K., S.M., A.V., D.T., A.I.A., S.C.R.W., J.C.W.B., D.J.L., and M.A.H.

## Acknowledgements

We thank the radiographers for their help with MRI scanning and all volunteers for their participation in the study.

