## Supplementary for "Test-retest reliability of sensorimotor activity measured with spinal cord fMRI"

### 1 Motor performance in each trial across task blocks

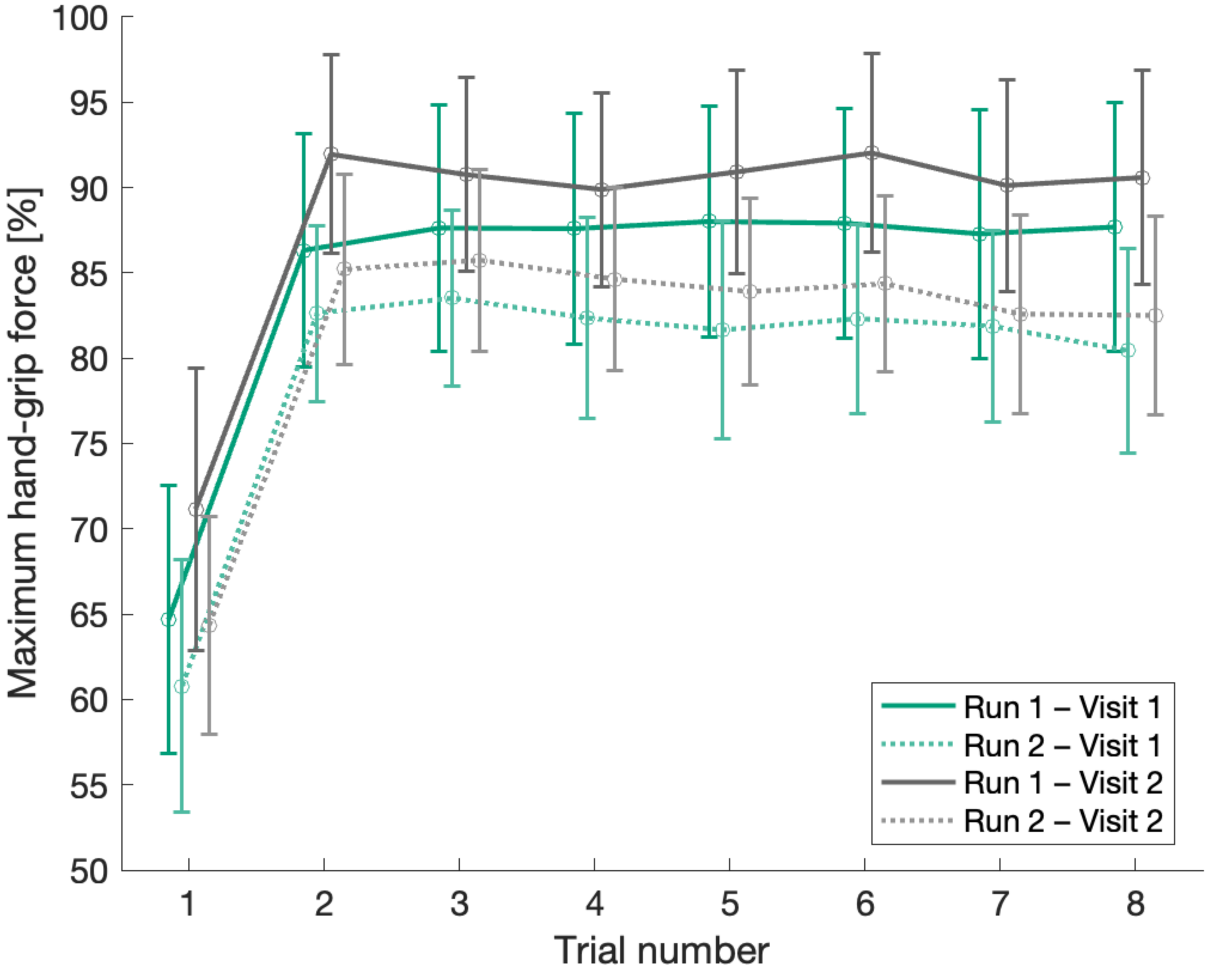

**Supplementary Figure 1.** Mean (± SD) hand-grip force exerted in each trial across the task blocks in each run of the task. The values are expressed as percentage of individually thresholded grip force.

### 2 Grip force-adjusted motor activation averaged across all four task runs

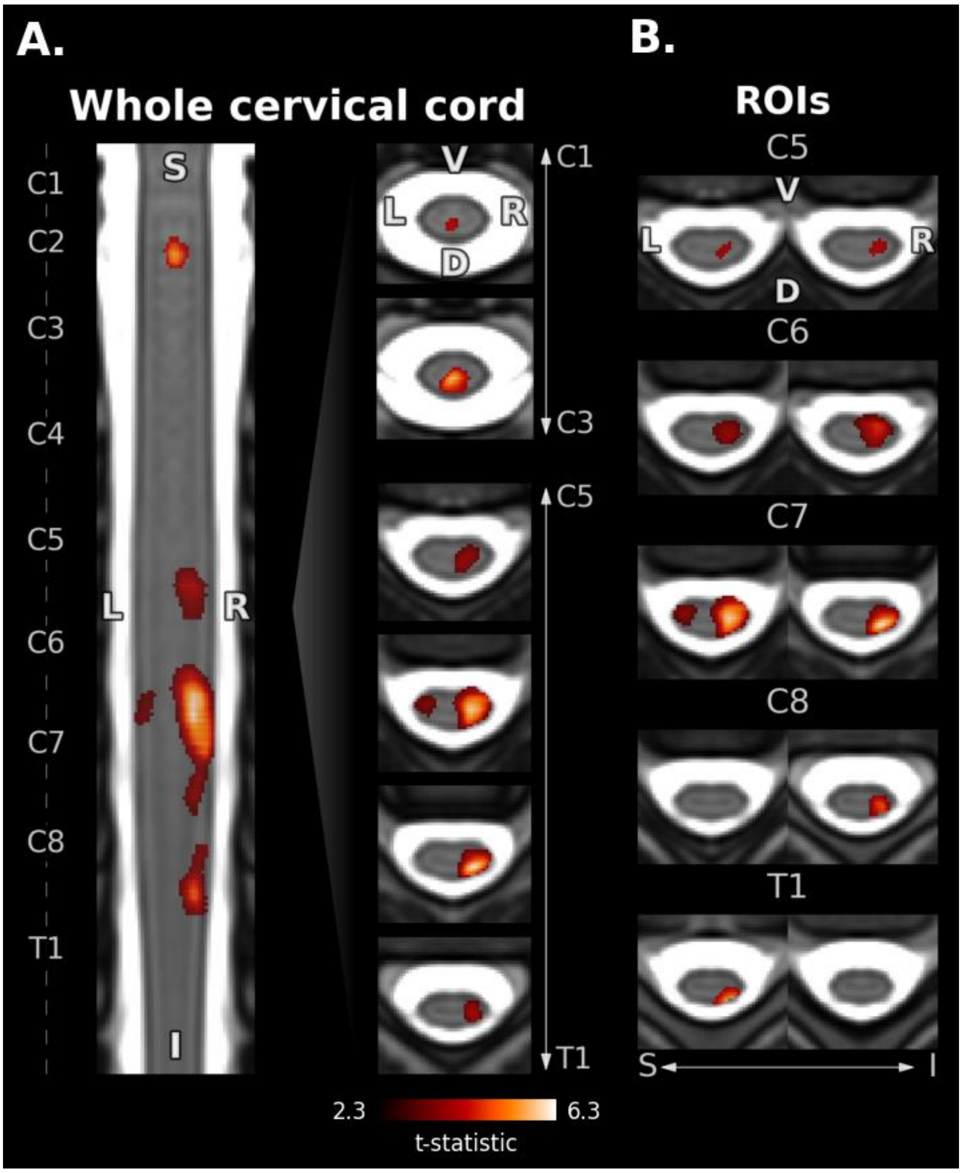

**Supplementary Figure 2.** Group-level grip force-adjusted motor activation averaged across all task runs. Statistically significant t-statistics are shown ($p_{FWE} < 0.05$). Labels denote spinal segmental levels. **A.** Activation obtained from whole cervical cord analysis shown on one representative coronal slice and representative axial slices from spinal segmental levels C1-C3 and C5-T1. **B.** Activation obtained from ROI analysis at each spinal segmental level C5-T1.

D = dorsal, I = inferior, L = left, n.s. = non-significant, R = right, ROI = region of interest, S = superior, V = ventral.

### 3 Grip force-adjusted motor activation in each task run

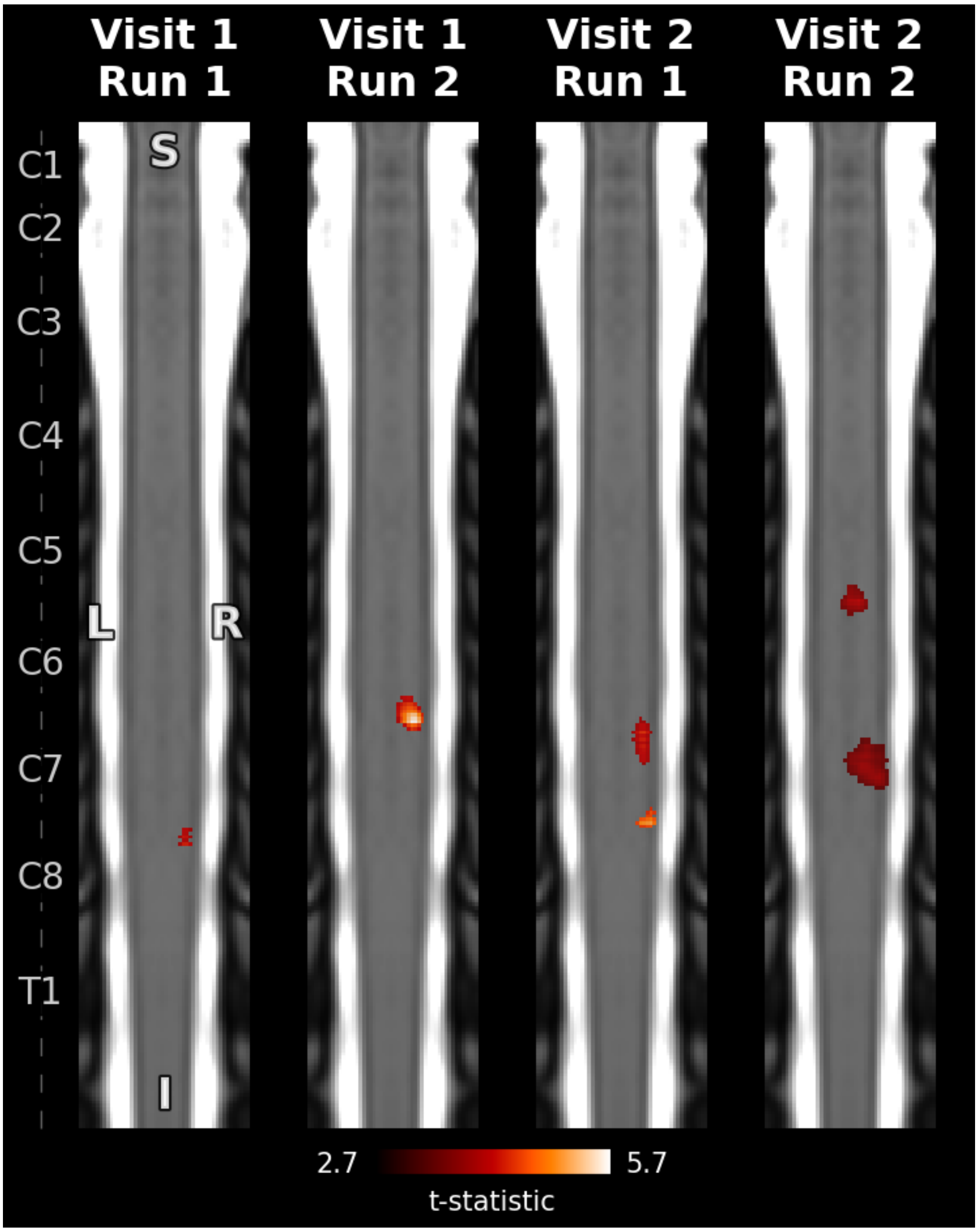

**Supplementary Figure 3.** Grip force-adjusted motor activation in each run of the task assessed within the whole cervical cord. One representative coronal slice per task run is presented. Statistically significant t-statistics are shown ($p_{FWE} < 0.05$). Left axis denotes spinal segmental levels.
I = inferior, L = left, R = right, S = superior.

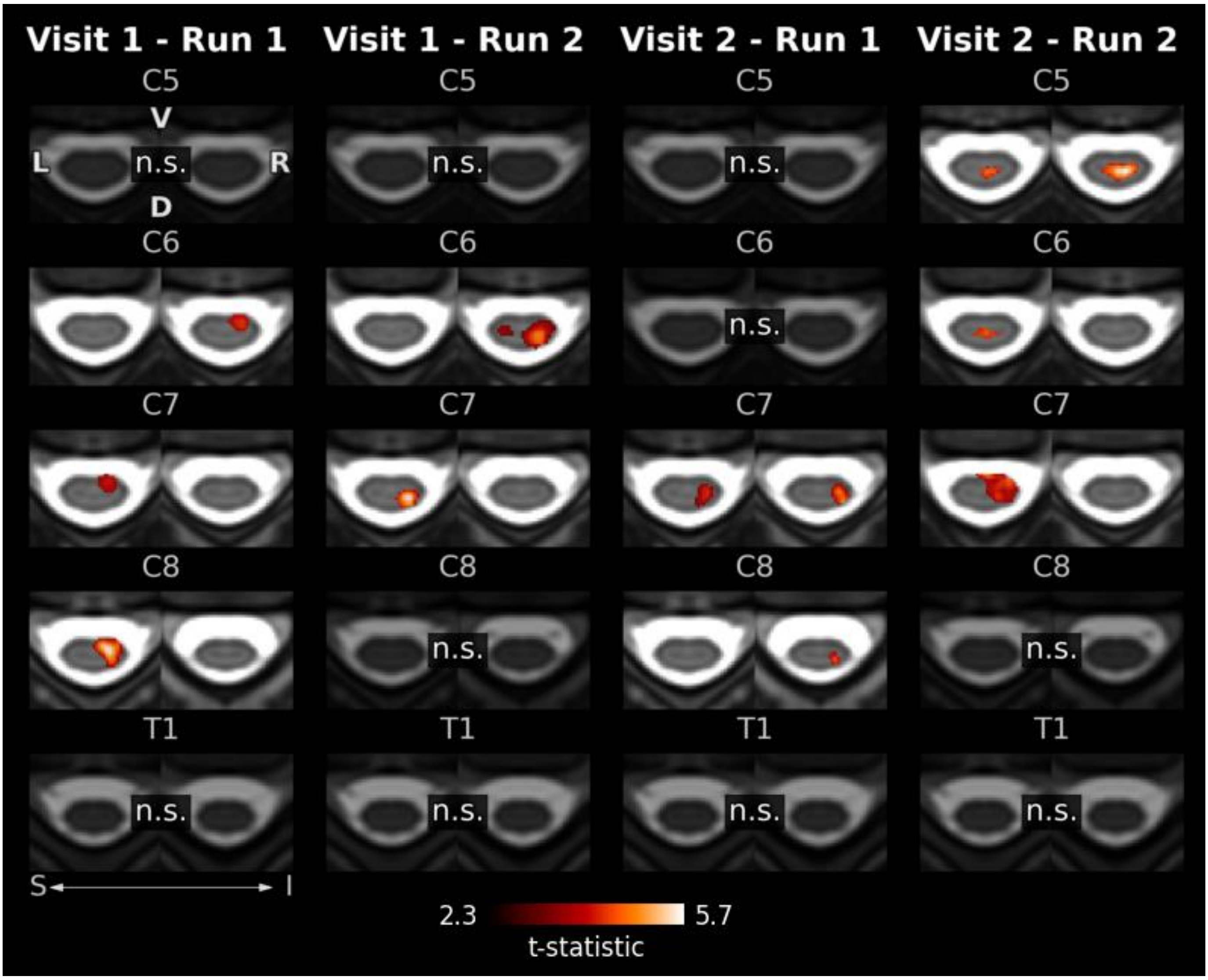

**Supplementary Figure 4.** Grip force-adjusted group-level motor activation in each run of the task assessed within each spinal segmental level ROI (C5-T1). Statistically significant t-statistics are shown ($p_{FWE} < 0.05$).
D = dorsal, I = inferior, L = left, n.s. = non-significant, R = right, ROI = region of interest, S = superior, V = ventral.

### 4 Impact of data quantity on grip force-adjusted motor activation

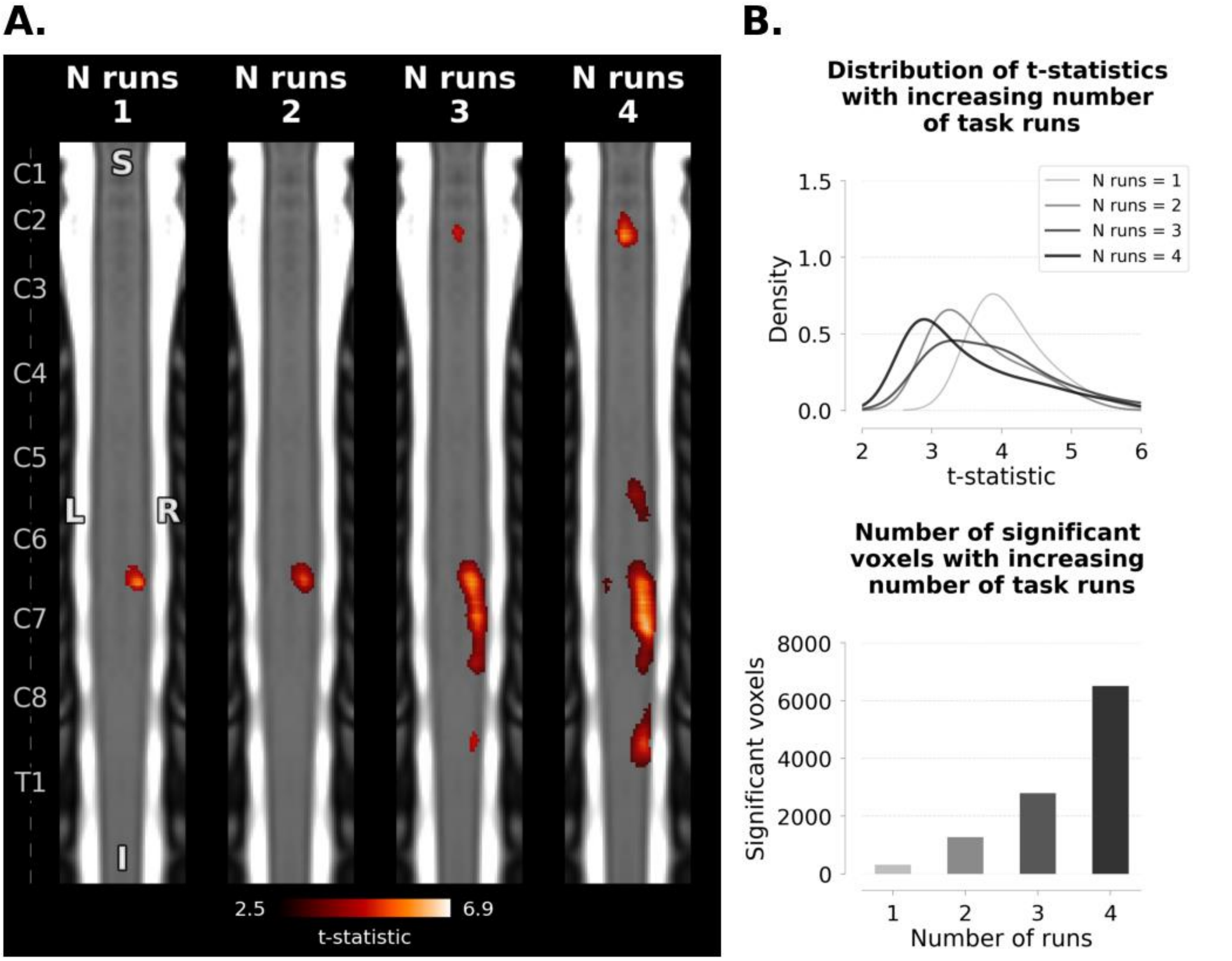

**Supplementary Figure 5.** Investigation of the impact of increasing the number of task runs included in analysis on group-level grip force-adjusted motor activation across the cervical spinal cord ($p_{FWE} < 0.05$). **A.** Group-level motor activation maps. One representative coronal slice per group-level is presented. Left axis denotes spinal segmental levels. **B.** Graphs showing the density of t-statistics (top) and number of active voxels (bottom) across the cord with increasing the number of task runs. I = inferior, L = left, N = number, R = right, S = superior.

### 5 Test-retest reliability of grip force-adjusted motor activation (parametric modulation)

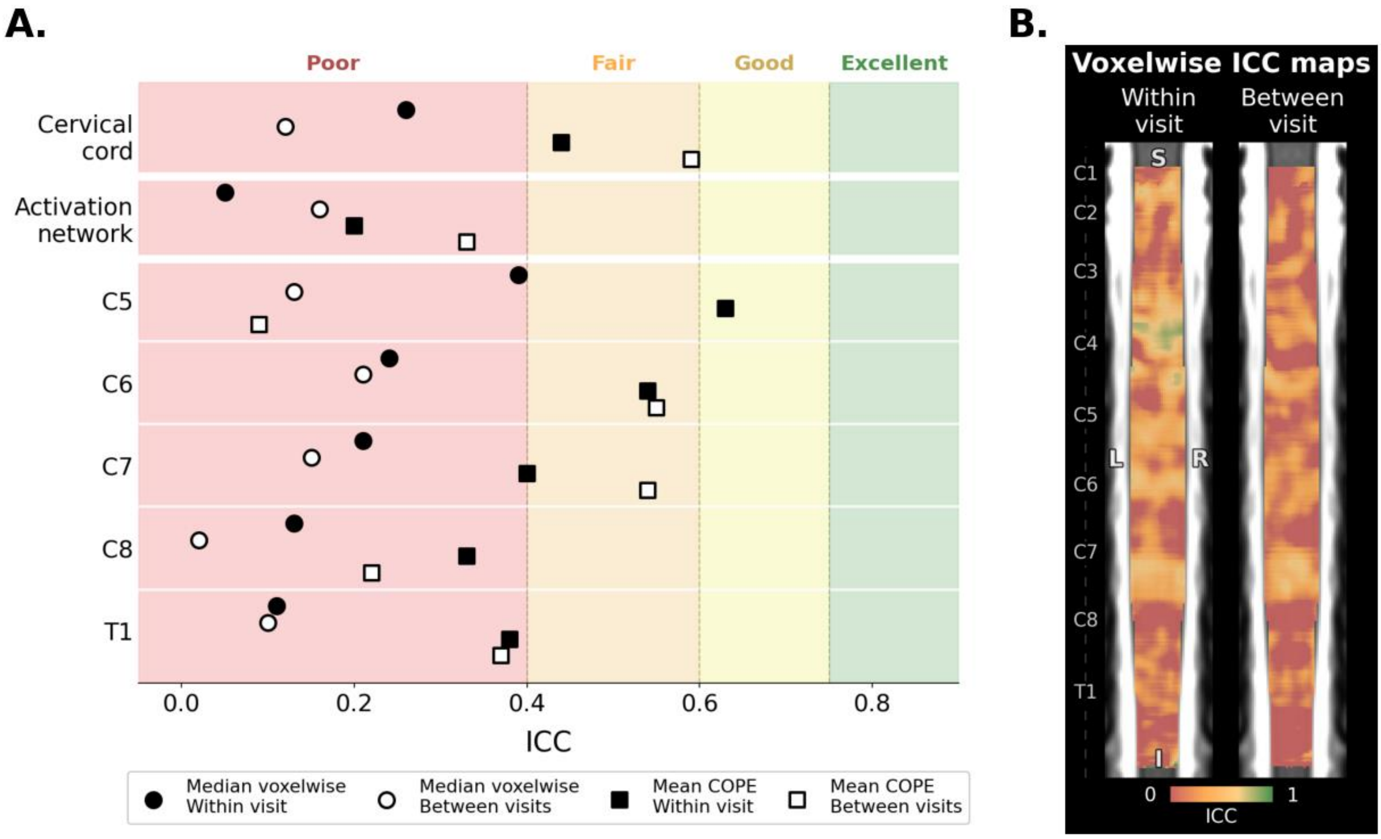

**Supplementary Figure 6.** ICC estimates for within- and between-visit reliability of grip force-adjusted motor activation. **A.** Forest plot showing median ICC for voxelwise assessments and ICC of mean COPE across the cervical spinal cord, the motor activation network, and spinal segmental levels C5-T1. **B.** Statistical maps showing ICC distribution for individual voxels within the cervical cord for within- and between-visit reliability assessments.
COPE = contrast of parameter estimates, ICC = intraclass correlation coefficient, SE = standard error.
